# Reduction of right-hemispheric auditory cortex activity in response to speech in more experienced paediatric cochlear implant users

**DOI:** 10.1101/2024.04.08.588535

**Authors:** Björn Kropf, Madhuri Sharma Rao, Alexander Gutschalk, Martin Andermann, Mark Praetorius, André Rupp, Kurt Steinmetzger

## Abstract

In case of severe hearing loss early in life or congenital deafness, cochlear implants (CIs) represent the method of choice to restore hearing and enable language acquisition. While speech intelligibility has been shown to improve during the first year after implantation to then reach a plateau, the underlying neuroplastic changes are poorly understood. Here, we longitudinally compared the cortical processing of speech stimuli in a case-control design with two groups of pre-lingually deafened CI users (4.4 vs. 25.8 months of CI experience) and an age-matched control group with normal hearing (NH; mean group ages ∼9 years). In two experiments, participants were presented with running speech and vowel sequences while fNIRS and EEG data were obtained simultaneously. Despite trends in this direction, cortical activity did not increase significantly with more CI experience and did not approach the higher levels observed in the NH controls. However, in the speech experiment, the less experienced CI group showed an abnormal shift of activity to the right hemisphere not observed in the other two groups. These results hence imply that adaptation to CI-based hearing is not characterised by a gradual increase of activity in left-hemispheric language network, but a reduction of abnormal activity elsewhere.

## 1. Introduction

Cochlear implants (CI) can enable hearing in patients with profound sensorineural hearing loss by directly stimulating the auditory nerve (Carlyon & Goehring, 2021; Macherey & Carlyon, 2014; Wilson & Dorman, 2008). CIs have proven successful in restoring hearing, allowing post-lingually deafened adults (Cullington & Zeng, 2008; Friesen et al., 2001; Stickney et al., 2004), and even pre-lingually deafened children (Litovsky et al., 2004; S. D. Sharma et al., 2020), to understand speech. However, a major limitation of CIs is that they provide limited access to pitch cues (Green et al., 2005; Oxenham, 2018; Steinmetzger & Rosen, 2018), resulting in impaired prosody perception (Meister et al., 2009; Nakata et al., 2012) and major difficulties in understanding speech in the presence of background noise (Dorman et al., 1998; Fu & Nogaki, 2005; Kwon et al., 2012). In adult CI users, speech intelligibility usually plateaus about 1 year after implantation and performance improvements are most pronounced during the first few months of CI use (Kral et al., 2019; Wilson & Dorman, 2008). However, the underlying neuroplastic changes following implantation in pre- as well as post-lingually deafened CI users are poorly understood and have to date mainly been investigated using animal models (Glennon et al., 2020; Kral et al., 2019).

Especially in paediatric CI users that have not yet acquired language, reliable behavioural data to assess their hearing are difficult to obtain. Consequently, measures of cortical activity that allow for an objective evaluation of their hearing have received considerable attention. Many of the resulting studies have used EEG, even though electrical artefacts caused by the CIs compromise the data quality (Gilley et al., 2006; Viola et al., 2011). Auditory ERPs recorded from paediatric subjects usually exhibit a prominent P1 whose latency decreases with age (Ponton et al., 2002; A. Sharma et al., 1997). A discernible P1 has been observed after about 6 months of CI use in young children (Ni et al., 2021), but compared to normal-hearing (NH) controls, the P1 in paediatric CI users is smaller and delayed (Gilley et al., 2008; Ponton et al., 1996; A. Sharma et al., 2002). A similar pattern has been reported for the MMN (Ni et al., 2021; Vavatzanidis et al., 2015), whereas the N400, which is thought to reflect semantic rather than acoustic processing of speech, was only observed after more than 1 year in infant CI users (Vavatzanidis et al., 2018). Furthermore, analyses of the latency and the cortical generators of the P1 have shown that adaptive neuroplasticity in pre-lingually deafened children appears to be maximal if they were implanted until about 3.5 years of age (‘critical period’) and declines markedly after the end of the so-called sensitive period at about 7 years (Gilley et al., 2008; A. Sharma et al., 2002). Moreover, permanently increased P1 amplitudes in the contralateral auditory cortex after more than 1.5 years of unilateral CI use suggest that pre-lingually deafened children should be provided with bilateral CIs in quick succession (Gordon et al., 2013). A common feature of these EEG studies is that they used typical ERP paradigms characterised by short, repetitive stimuli rather than natural speech.

In contrast, a second line of research used continuous speech to investigate neuroplastic changes in paediatric CI users. Most of these studies used fNIRS, which is considered ideal to study the spatial distribution of cortical activity in this population as the implants do not interfere with the measurements and due to its ease of use (Bortfeld, 2019; Pinti et al., 2018; Saliba et al., 2016). In an early combined fNIRS and fMRI study, speech-evoked activity was seen in most tested children directly after CI switch on as well as after 4 months, but neither topographies of cortical activity nor comparisons with the NH control group were provided making it difficult to draw conclusions from these results (Sevy et al., 2010). More recent fNIRS studies reported bilateral auditory cortex activity in response to speech, with no differences in activity between pre-lingually deafened children with several years of CI experience and NH controls (Mushtaq et al., 2020) or even larger responses in the CI group (Zhou et al., 2023), and similar activation patterns in infant CI users directly after the devices were switched on and 1.5 months later (Wang et al., 2022). Moreover, in a longitudinal PET study (Petersen et al., 2013), speech elicited left-lateralised auditory cortex activity in post-lingually deafened adult CI users within the first 6 months after switch-on, while no such effect was evident in a small group of 4 pre- lingually deafened adult CI users. Thus, the few available studies suggest no left-lateralisation of speech-evoked cortical activity in pre-lingually deafened paediatric CI users, contrary to what was found in NH newborns and infants (Dehaene-Lambertz et al., 2002; Minagawa-Kawai et al., 2011; Pena et al., 2003; H. Sato et al., 2012) as well as children (Berl et al., 2014; Lawrence et al., 2021). However, none of the respective fNIRS studies (Mushtaq et al., 2020; Sevy et al., 2010; Wang et al., 2022; Zhou et al., 2023) provided evidence for diminished auditory cortex activity in CI users relative to NH controls, a typical finding when contrasting these listener groups using ERP as well as neuroimaging techniques (e.g., Coez et al., 2008; Gilley et al., 2008; Sandmann et al., 2009; Steinmetzger et al., 2022a). The absence of this effect thus raises the question whether fNIRS is at all suitable for reliably detecting auditory cortex activity in paediatric subjects. Furthermore, changes in speech-evoked cortical activity were so far only assessed during the first few months after implantation (Petersen et al., 2013; Sevy et al., 2010; Wang et al., 2022) and it thus remains unclear if left-lateralised activity in pre-lingually deafened paediatric CI users might emerge with more CI experience. Additionally, only two fNIRS studies to date investigated the cortical processing of prosodic features in paediatric CI users (Chen et al., 2022; Wang et al., 2021) and neither of them reported the right-lateralisation of activity in response to prosodic modulations typically observed in NH infants (Homae et al., 2006; Homae et al., 2012; Telkemeyer et al., 2011) and children (Wartenburger et al., 2007). Yet, both of them were cross-sectional studies and provide no information regarding neuroplastic changes following implantation.

Here, we employed a longitudinal design in which pre-lingually deafened paediatric CI users (mean age ∼9 years) were tested during and after their first year of CI use and compared them to an age-matched NH control group. To close the gap between the electrophysiological and neuroimaging studies summarised above, fNIRS and EEG data were obtained simultaneously while the participants listened to continuous speech as well as to vowel sequences with different prosodic properties. The concurrent use of both methods also enabled the mutual validation of the results. The EEG data recorded in response to running speech were used to derive temporal response functions (TRFs), which represent ERP-like auditory evoked responses to continuous stimuli (Crosse et al., 2016; Lalor & Foxe, 2010). This model-based approach enabled the direct comparison of the EEG and fNIRS data obtained in the speech experiment. In the subsequent vowel experiment, continuous vowel sequences in which the prosodic features were either fixed or variable between the individual vowels were contrasted. The same stimulus paradigm was used in earlier combined fNIRS and EEG studies with adult NH listeners (Steinmetzger et al., 2022a) and post-lingually deafened adult CI users (Steinmetzger et al., 2022b). Both of these studies provided evidence for a right lateralisation of the processing of vowel sequences with variable prosodic features, in line with studies that have investigated prosody perception in paediatric NH subjects (Homae et al., 2006; Homae et al., 2012; Telkemeyer et al., 2011; Wartenburger et al., 2007).

## 2. Material and methods

### 2.1. Participants

Thirteen pre-lingually deafened paediatric CI users and 13 age-matched NH control participants took part in this case-control study. Detailed information regarding the participants is provided in Table 1. The participants or their legal guardians gave written consent prior to the experiment. The study was approved by the local research ethics committee (Medical Faculty, University of Heidelberg) and was conducted in accordance with the Declaration of Helsinki.

The data were subdivided into three groups to assess neural plasticity effects following CI use and to compare CI-based electrical hearing to normal acoustic hearing. The grouping and the measurement time points for each CI subject are shown in Fig. 1A. Group *CI – T1* included datasets recorded during the first year of CI use. This group contained 17 datasets from 8 different CI subjects, 6 of which were tested repeatedly [mean age (years.months) = 8.4, SD = 5.7]. Group *CI – T2* consisted of datasets recorded after the first year of CI use and comprised 12 datasets from 12 different participants, 7 of which were also part of the first group (mean age = 9.3, SD = 5.9). Although the deafness periods prior to receiving their CIs differed widely across subjects, there was no significant difference between the datasets in the CI – T1 and CI – T2 groups (means = 7.11/7.2, SDs = 5.10/6.3; *t*_(27)_ = 0.35, *p* = 0.732). The *NH controls* group included 13 subjects (mean age = 9.2, SD = 5.8).

**Figure 1.**
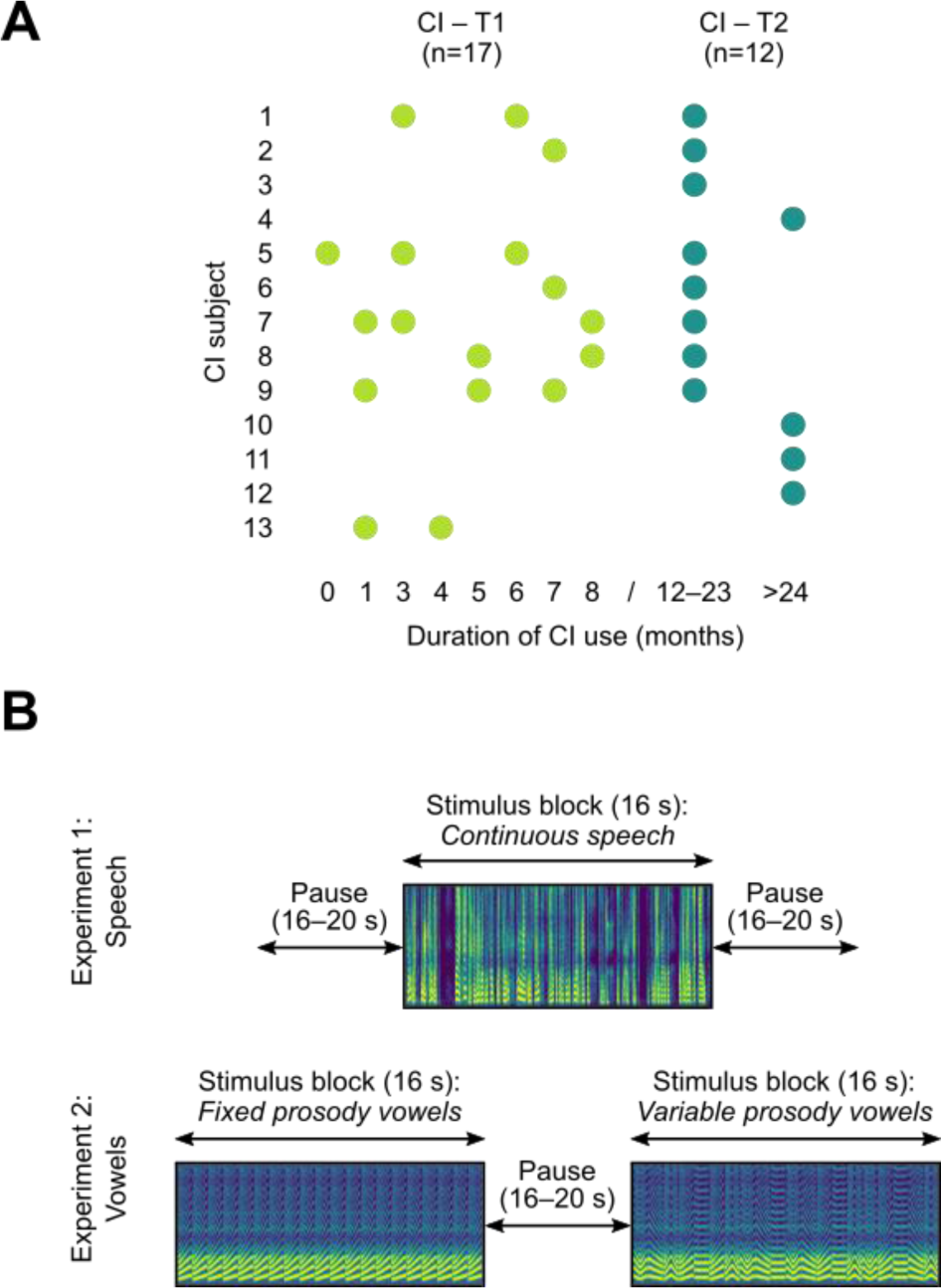
Experimental design and stimuli. A) The datasets obtained from the CI users were divided into those collected during (CI – T1, mean duration of CI use = 4.4 months) and after the first year of CI use (CI – T2, mean duration = 25.8). B) Schematic illustration of the block design used in the speech (top row) and vowel (bottom row) experiments. Narrow-band spectrograms of the stimuli are shown to emphasise spectral differences between the FIXED and VARIABLE PROSODY conditions in the vowel experiment.

CI subject 1 had received a second CI shortly before the 3^rd^ testing session, but the measurement time points are given relative to the first CI. For CI subject 5, in contrast, the time points are given relative to the second CI, received about 1 year after the first one. Here, it was assumed that the adaptation to the first CI was almost complete, and that the subject’s young age and relatively short deafness period would also result in a large degree of neural plasticity in response to the second CI. The other two bilaterally implanted subjects (nos. 10 & 13) received their implants simultaneously.

| Subject | Age<sup>1</sup><br>(years.months) | Sex | CI<br>ear/s | Deafness<br>duration | Aetiology of deafness | Implant & processor type / strategy | Words correct<sup>2</sup> | Age NH<br>control |
| --- | --- | --- | --- | --- | --- | --- | --- | --- |
| 1 | 22.6 | f | both | 21.6 | Connexin 26, 30delG-mutation | CI622 & KANSO / ACE, bilaterally | 3, 6, 12 months: 60%, 60%, 80% | 23.3 |
| 2 | 2.5 | f | l | 1.5 | Cochlear nerve aplasia | FLEX28 & SONNET2 / FS4 | - | 2.6 |
| 3 | 18.2 | f | r | 16.3 | Recurrent otitis media | FLEX28 & SONNET / FS4-p | 12 months: 65% | 18.4 |
| 4<sup>3</sup> | 4.8 | f | r | 2.6 | Large aqueduct syndrome | CI522 & CP1000 / ACE | 24 months: 70% (Göttinger II) | 5.0 |
| 5<sup>4</sup> | 7.1 | f | both | 4.11 | Unknown, probably congenital | CI622 & CP1000 / ACE, bilaterally | 8 months: 60% (Göttinger I) | 6.11 |
| 6<sup>4</sup> | 7.10 | m | r | 6.10 | Unknown | FLEX28 & SONNET / FS4 | 12 months: 35% | 7.7 |
| 7<sup>4</sup> | 9.3 | f | l | 8.3 | Unknown | FLEX28 & SONNET2 / FS4 | 6, 12 months: 40% (Mainzer II), 90% (Göttinger II) | 8.0 |
| 8 | 2.7 | m | r | 1.6 | Hyperbilirubinemia | FLEX28 & SONNET2 / FS4 | 9 months: 60% (Mainzer I) | 3.7 |
| 9 | 10.7 | m | r | 9.6 | Mondini, widened vestibular aqueduct | FLEX26 & SONNET2 / FS4 | 3, 6, 9, 12 months: 60%, 30%, 40% (Mainzer I), 30% (Mainzer II) | 10.0 |
| 10 | 10.6 | m | both | 0.10 | Unknown | CI522 & CP1000 / ACE, bilaterally | - | 11.1 |
| 11<sup>3</sup> | 8.7 | m | r | 6.3 | Unknown | CI522 & CP1000 / ACE | - | 9.6 |
| 12 | 8.0 | m | r | 5.10 | Icterus of the newborn | CI622 & CP1000 / ACE | 12 months: 50% | 7.3 |
| 13<sup>4</sup> | 6.4 | m | both | 6.0 | Unknown | FLEX28 & SONNET2 / FS4, bilaterally | - | 6.10 |

<sup>1</sup> Age at last test session. <sup>2</sup> Words correct scores were determined with the Freiburg monosyllabic speech intelligibility test (Hahlbrock, 1953) at a presentation level of 65 dB SPL, unless noted otherwise. <sup>3</sup> Hearing aid in contralateral ear taken off for testing. <sup>4</sup> No German native speakers.

### 2.2. Stimuli

In the first part of the experiment, participants were presented with excerpts from a recording of the fairy tale “*Tischlein deck Dich”*, read in an animated, child-friendly manner by an adult female German talker. The 16-s excerpts were tapered on and off using 50-ms Hann windows, low-pass filtered at 3.5 kHz (zero-phase-shift 1^st^-order Butterworth), and normalised to a common root-mean-square level.

The vowel stimuli used in the second part of the experiment were identical to those employed in previous fNIRS-EEG experiments with adult NH listeners (Steinmetzger et al., 2022a) and adult CI users with single-sided deafness (Steinmetzger et al., 2022b), where the stimulus construction is described in more detail. The stimulus materials were recordings of the German vowels /a/, /e/, /i/, /o/, and /u/ spoken by an adult male German talker in an anechoic room. Each vowel was limited to a length of 800 ms using a 50-ms Hann-windowed offset ramp, and re-synthesised with a range of prosodic contours (*flat*, *rising straight*, *falling straight*, *rising curved*, and *falling curved*) with the STRAIGHT vocoder software (Kawahara & Irino, 2005). For the non-flat contours, the F0 increased or decreased by a perfect fifth relative to the mean F0 of the contour.

### 2.3. Experimental design and procedure

The first part of the experiment consisted of 10 blocks of continuous speech in fixed order, interspersed with pauses (Fig. 1B). Hence, this part of the experiment had a duration of about 6 min. The short duration was to ensure that even the youngest participants were able to complete this part of the experiment as well as a portion of the subsequent vowel experiment.

The design of the second part of the experiment was identical to previous studies (Steinmetzger et al., 2022a; Steinmetzger et al., 2022b). The individual vowels were presented in continuous blocks of 20, alternating with pauses (Fig. 1B). The experiment consisted of two conditions: In the FIXED PROSODY condition, all vowels within a block had the same prosodic contour (*flat*, *rising straight*, *falling straight*, *rising curved*, or *falling curved*). In the VARIABLE PROSODY condition, the contours varied between the vowels within each block (*rising*, *falling*, *straight*, *curved*, or a *mixture of all five contour types*). The five different contour types were intended to represent a set of typical, easily distinguishable prosodic contours found across languages. In total, the second part of the experiment comprised 100 blocks, 50 per condition, presented in random order, amounting to a duration of about 57 mins. As the EEG data were analysed relative to the onset of the individual vowels, this design resulted in 2000 trials. The randomisation procedure was set so that the number of blocks per condition was approximately the same regardless of when the experiment was terminated, as the younger children were not expected to complete the experiment. On average, the participants completed 79.9 blocks per session (CI data sets = 82.0, min = 22; NH data sets = 75.3; min = 24).

Participants were tested using free-field acoustic stimulation, for which the stimuli were converted with 24-bit resolution at a sampling rate of 48 kHz using an ADI-8 DS sound card (RME, Haimhausen, Germany) and presented via an Adam A7x speaker (Adam Audio, Berlin, Germany). The speaker was placed directly in front of the listener, ∼1.5 m away and at ear level. The presentation level was set to 70 dB SPL using a sound level meter (Brüel & Kjær, type 2235; Nærum, Denmark) located at the position of the subject’s head. The experiment took place in an acoustically and electrically shielded room. Depending on age, height, and personal preference, the participants either sat in a comfortable reclining chair, a child car seat, or on the lap of their accompanying legal guardian during data acquisition. To minimise ambient light from interfering with the fNIRS recordings, the participants wore an overcap and the room light was dimmed to the lowest level. There was no behavioural task, but pauses were inserted about every 10 mins to ensure the vigilance of the subjects.

### 2.4. fNIRS recording and analysis

fNIRS signals were recorded with a continuous-wave NIRScout 16×16 system (NIRx Medizintechnik, Berlin, Germany) at a sampling rate of 7.8125 Hz. The source optodes emitted infrared light with wavelengths of 760 and 850 nm. Eight source optodes and eight detector optodes were placed symmetrically over each hemisphere by mounting them on a custom EEG cap (EasyCap, Herrsching, Germany). The chosen optode layout was devised to optimally cover the auditory cortex and adjacent areas. This layout resulted in 22 measurement channels per hemisphere, of which 20 had a standard source-to-detector distance of about 30 mm, while the remaining 2 had a shorter spacing of about 14 mm. The optode and reference positions for two subjects (CI 1 & CI 9) were digitised with a Polhemus 3SPACE ISOTRAK II system (Colchester, VT, USA) before the recordings. These two digitisations, which showed a close spatial agreement (Suppl. Fig. 1.), served as templates for all adult (CI 1) and minor-aged subjects (CI 9), respectively.

The raw data from the speech and vowel experiments were separately pre-processed using the HOMER2 toolbox (version 2.8; Huppert et al. 2009) and custom MATLAB code. The raw light intensity signals were first converted to optical density values and then corrected for motion artefacts. A kurtosis-based wavelet algorithm with a threshold value of 3.3 was used to identify and correct motion artefacts (Chiarelli et al., 2015). Measurement channels with poor signal quality were then identified by computing their scalp coupling index (SCI; Pollonini et al. 2014) and excluded from further analysis if the SCI value was smaller than 0.5. As we did not pre-select the subjects, the sample included several participants with long, dark hair. Thus, a lower SCI threshold than the one suggested by Pollonini and colleagues (0.75) was used to limit data loss. A maximum of 12 channels per subject and experiment were excluded (speech experiment: mean CI data sets = 3.0, mean NH = 0.3; vowel experiment: mean CI = 3.1, mean NH = 1.2), except for one subject (CI 7) with very thick dark hair (max 20 channels across experiments and sessions, mean = 14.9). Next, the motion-corrected signals of the remaining channels were band-pass filtered between 0.01–0.5 Hz to isolate the task-related neural activity, and subsequently converted to concentration values based on the modified Beer-Lambert law (Scholkmann et al., 2014). The differential path length factors required for the conversion were determined based on the wavelengths of the light and the age of the subject (Scholkmann & Wolf, 2013).

The pre-processed data were further processed, statistically evaluated and topographically visualised using a customised version of SPM-fNIRS (version r3; Tak et al. 2016). The optode positions were first transformed from subject space to MNI space, after which they were probabilistically rendered onto the ICBM-152 cortical template surface (Singh et al., 2005). The pre-processed signals were then temporally smoothed using the shape of the canonical haemodynamic response function (HRF, ‘pre-colouring’) to avoid autocorrelation issues (Worsley & Friston, 1995). Furthermore, the signals from the four short channels were subjected to a principal component analysis, using the first component as a nuisance regressor to remove the so-called global scalp-haemodynamic component (T. Sato et al., 2016), i.e., the superficial signal component which is thought to not reflect any cortical activity. In the resulting topographical plots, the short channels are thus omitted.

In contrast to our earlier fNIRS studies with adult participants (Steinmetzger et al., 2022a; Steinmetzger et al., 2022b; Steinmetzger et al., 2020), the statistical evaluation of the data was based on mean amplitude measures rather than statistical parametric mapping (SPM). This approach was chosen as the SPM-based topographies of functional activity in the speech experiment were very similar across groups despite markedly different waveform morphologies, indicating a limited sensitivity for detecting differences between the paediatric samples in the current study. Hence, the mean amplitudes from 0–32 s past stimulus onset were extracted for each subject and experimental condition to enable a model-free assessment of the functional activations. The HRFs were baseline corrected by subtracting the mean amplitude from −2–0 s before stimulus onset from each sample point. In the HRF plots, the waveforms are shown both after the pre-processing (‘Total HRF’) as well as after regressing out the contribution of the short channels and pre-colouring the signals (‘Cortical HRF’), to illustrate the effect of removing the non-cortical signal component. Group-level statistics for each long channel were computed based on the subject-level amplitudes of the cortical HRFs using one-sided *t*-tests.

For comparison, SPM-based analyses are provided as supplementary material (Suppl. Fig. 2). Here, the data of the individual subjects were modelled by convolving the continuous signals obtained from each long channel with 16-s canonical SPM double-gamma functions representing the stimulus blocks. The oxygenated (HbO) and de-oxygenated (HbR) haemoglobin data were modelled with positive and negative HRFs, respectively. To allow the time course of the measured concentration changes to vary slightly, the temporal and spatial derivatives of the canonical HRF were included as additional regressors. After estimating the HbO and HbR general linear models (GLMs) for each subject, contrast vectors were defined to assess the functional activations. When comparing the activity across groups or conditions, the regressors of interest were set to 1 and −1, respectively, while the regressors representing the derivatives and the global scalp component were set to 0 to statistically control for their effects. Likewise, when evaluating the activity within groups or conditions against baseline, the respective regressors were set to 1, whereas all other regressors were set to 0. Group-level statistics for each long channel were computed based on the subject-level beta weights using one-sided *t*-tests.

One NH control subject (NH 11) was excluded from all fNIRS analyses because the HbR results in the speech experiment showed no activity in the auditory cortex, while the HbO amplitudes were overly negative (*z*-score of the mean across all channels = −2.74). Throughout the present study, we focused on the HbR data as these have proven to be a more reliable measure of auditory activity than HbO data, which often exhibited negative responses in our previous studies (Steinmetzger et al., 2022b; Steinmetzger et al., 2020). However, the HbO results are provided as supplementary material (Suppl. Fig. 3).

### 2.5. EEG recording and analysis

Continuous EEG signals were recorded using a BrainVision actiCHamp system (Brain Products, Gilching, Germany). Depending on the head size, a 64-(n = 6) or 32-channel (n = 30) setup was employed. However, for the 64-channel recordings, only the first 32 channels were considered in the analyses to ensure consistency. CI-compatible custom EEG caps with holes for the transmitter coils at electrode positions P7 and P8 were used. Apart from this deviation, the scalp electrodes were arranged according to the extended international 10-20 system. For the 64-channel recordings, 4 electrodes were placed around the eyes to record vertical and horizontal eye movements. The EEG data were recorded with an initial sampling rate of 500 Hz, an online anti-aliasing low-pass filter with a cut-off frequency of 140 Hz, and were referenced to the right mastoid. In some of the sessions, no EEG signals could be recorded as the participants were too restless for an adequate preparation of the measurements, resulting in group sizes of n = 15 (CI – T1), n = 10 (CI – T2), and n = 11 (NH controls).

The raw data were pre-processed offline using FieldTrip (version 20180924; Oostenveld et al. 2011) and custom MATLAB code. For the speech experiment, the continuous waveforms were first segmented into epochs ranging from −4.1–20.1 s relative to block onset. Next, the epochs were filtered using zero-phase shift Butterworth filters (high-pass cut-off 1 Hz, 3^rd^ order; low-pass cut-off 15 Hz, 4^th^ order). The epochs were then re-referenced to electrode Cz or an adjacent midline channel in case of bad signal quality and down-sampled to 250 Hz. After visually identifying and excluding bad channels (mean CI data sets = 5.9 channels, max = 15; mean NH = 5.5, max = 16; see Suppl. Fig. 4 for scalp distribution), the data were decomposed into 20 principal components to identify and reject eye artefacts. For about half the data sets (19/36), the principal component analysis was performed a second time to detect and eliminate CI-related and other artefactual signal components. Bad channels were then interpolated using the weighted average of the neighbouring channels and three fringe electrodes (TP9, TP10, and O1) that were identified as bad channels in more than 10 data sets were removed. Finally, the data were re-referenced to the average of the remaining channels.

For the vowel experiment, epochs ranging from −0.3–0.9 s relative to vowel onset were extracted and linear trends as well as the DC component were removed by subtracting a 1st-order polynomial instead of high-pass filtering. After excluding bad channels (mean CI data sets = 6.9 channels, max = 16; mean NH = 5.1, max = 16) and eye electrodes, epochs in which the amplitudes between −0.2–0.8 s around vowel onset exceeded ±125 µV or the z-transformed amplitudes differed by more than 15 standard deviations from the mean of all channels were excluded from further processing. In three data sets, the rejection thresholds were increased to ±250 µV to ensure enough trials. On average, 91% of the trials (1569/1729) passed the rejection procedure for the CI data sets and 88% for the NH data sets (1373/1555). All other pre-processing steps were the same as in the speech experiment.

The pre-processed data from the speech experiment were then used to derive temporal response functions (TRFs) to the envelope of continuous speech. Using the mTRF toolbox (version 2.3; Crosse et al., 2016), the broadband envelope of the speech blocks was extracted at a sampling rate of 250 Hz by averaging the squared signal amplitude at regular intervals, taking the square root, and logarithmically compressing the resulting RMS intensity. For each electrode, TRFs with lags between −0.15 and 0.65 s were computed by fitting a regularised linear regression model mapping the speech envelope to the observed neural signals. Both the individual stimulus envelopes and the EEG data of each channel were *z*-standardised before fitting the regression model. The ridge parameter that prevents over-fitting of the model was estimated for each individual data set by identifying the value that resulted in the lowest mean square error between stimulus envelope and EEG data. The estimated ridge parameters were then averaged across all trials and electrodes to determine the parameter value for a given data set. The set of tested ridge parameters comprised 100 log-spaced values between 10^0^ and 10^6^. The resulting TRFs for each electrode were again *z*-scored to control for amplitude differences between data sets due to different ridge parameter values. Lastly, the speech TRFs as well as the ERPs from the vowel experiment were baseline corrected by subtracting the mean amplitude for lags from −0.1–0 s or before stimulus onset, respectively.

## 3. Results

### 3.1. Speech experiment

The group-level fNIRS HbR topographies are shown in Fig. 2A, where the channel-wise functional activations in response to speech are depicted for each of the three groups. After applying a strict Bonferroni correction across all 40 measurement channels, significant activity in the CI – T1 group was confined to the right superior temporal cortex (STC; *p_(Bonf)_* < 0.05, ch# 28 & 34, averaged Cohen’s *d* = 1.25). In contrast, no significant activity was evident for the CI – T2 group. The NH control group, on the other hand, showed significant activity in the left STC (*p_(Bonf)_* < 0.05, ch# 12 & 16, *d* = 1.32).

**Figure 2.**
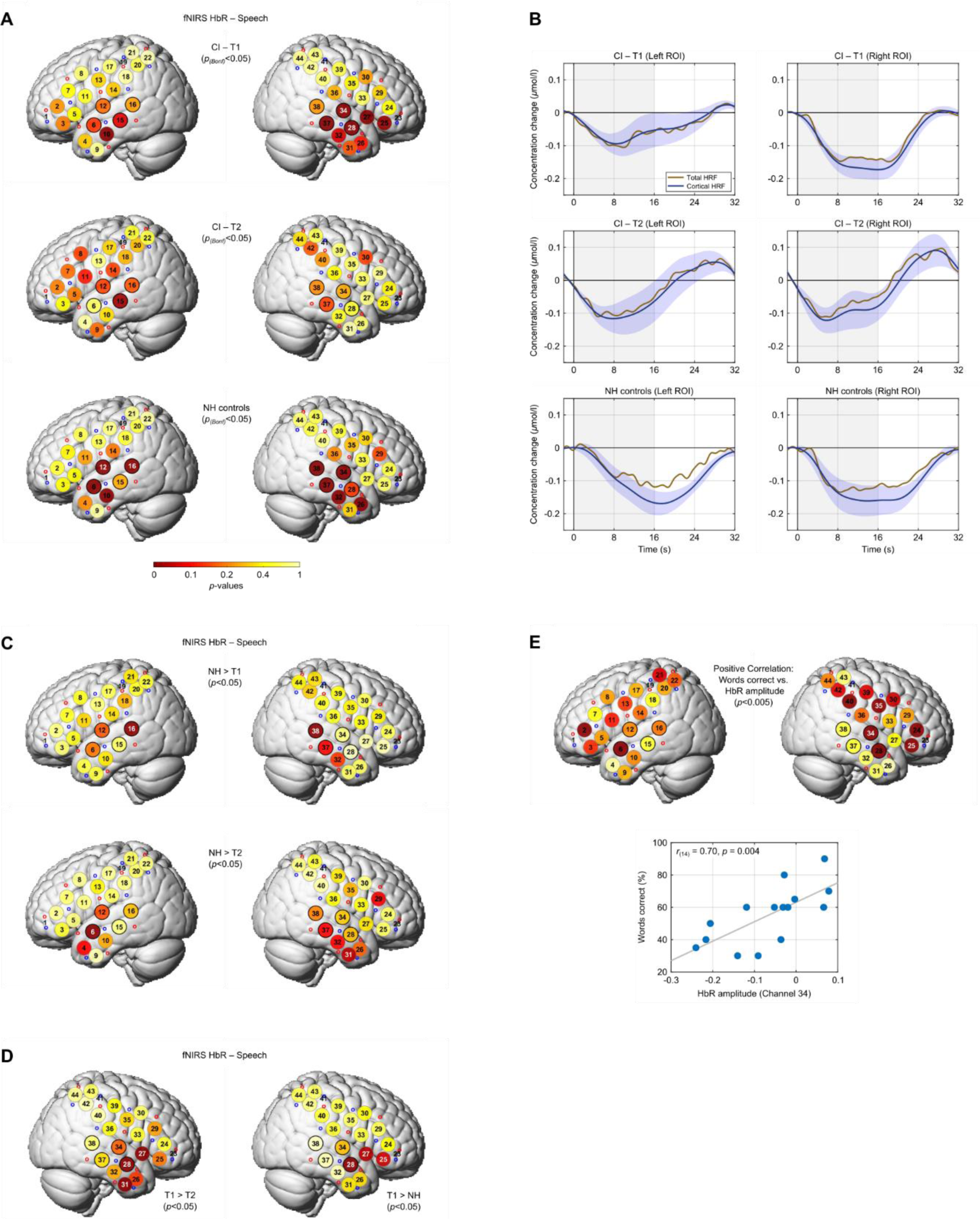
fNIRS results: speech experiment. A) fNIRS HbR topographies of functional activity in response to speech for each of the three experimental groups. Channels within the two auditory ROIs are outlined by black circles. White channel numbers indicate significant ROI channels. B) HRFs for each group and ROI. The shading indicates the standard error of the mean. C) Comparison of the topographies of functional activity across groups. D) Correlations of CI-based speech intelligibility and functional activity. The strongest positive correlation was observed for channel 34, as shown in the scatter plot. The words-correct scores are taken from Table 1.

To complement the topographical results, the corresponding HRFs are shown in Fig. 2B, separately for each group and hemisphere. The HRFs are shown for two pre-defined auditory regions of interest (ROIs), each comprising four channels in the mid superior temporal gyrus/sulcus (STG/STS; left: ch# 6, 12, 15 & 16; right: ch# 28, 34, 37 & 38), where speech typically evokes the largest activity (Alho et al., 2014). In line with the shapes of the HRFs, a right lateralisation was evident for the CI – T1 group (*t*_(15)_ = 2.43, *p* = 0.028, *d* = 0.57), while no hemispheric asymmetries were observed in the other two groups (*p* ≥ 0.701).

Next, the functional activation patterns were compared across groups (Fig. 2C). For the NH group, activity in the posterior portion of the auditory cortex was stronger compared to the CI – T1 group in both hemispheres (*p* < 0.05, ch# 16 & 38, *d* = 1.14). Furthermore, the NH control group showed greater activity in the left anterior STS, an area specifically activated by the processing of intelligible speech (Scott et al., 2000), than the CI – T2 group (*p* = 0.015, ch# 6, *d* = 0.99). Stronger activity in the NH group compared to the CI – T2 group was also observed in the right inferior temporal gyrus, outside the auditory ROI (*p* = 0.046, ch# 31, *d* = 0.73).

In turn, activity in the right anterior temporal lobe and inferior frontal gyrus was enhanced in the CI – T1 group compared to both the CI – T2 group (*p* < 0.05, ch# 27, 28 & 31, *d* = 1.35) and the NH controls (*p* < 0.05, ch# 25, 27 & 28, *d* = 0.81), as shown in Fig. 2D.

Additionally, we examined whether the CI subject’s speech intelligibility scores were reflected in the fNIRS HbR data obtained in response to running speech (Fig. 2E). For the 15 data sets for which clinical speech test data were acquired in close temporal proximity to the respective test sessions (Table 1), widespread positive correlations of speech scores and HbR amplitudes were evident in the right hemisphere. This effect was most pronounced in the right auditory cortex (*r_(14)_* = 0.70, *p* = 0.004, ch# 34), but also present in non-auditory areas (*p* < 0.005, ch# 25 & 35). Thus, better performance was associated with lower levels of cortical activity, as indicated by less negative HbR amplitudes.

The simultaneously recorded EEG data were analysed by estimating TRFs to the envelopes of the continuous speech blocks, which represent auditory evoked responses akin to typical ERPs. In a first step, the global field power (GFP) time courses of the grand-average TRFs were computed for each group (Fig. 3A). The GFP is defined as the variance across all electrodes and represents a reference-free measure of the overall strength of activity at a given point in time (Skrandies, 1990). In all three groups, the GFPs exhibited a two-peaked morphology, with a first peak around 70 ms and second one around 225 ms. The latter peak was substantially larger in the NH control group. Scalp maps comprising the latency window (125–325 ms) of the second peak are shown in Fig. 3B. While the largest mean amplitudes were observed in the fronto-central scalp region for the CI – T1 group, a shift towards left temporal areas is evident in the CI – T2 group. For the NH controls, activity was most pronounced in the left fronto-temporal scalp region. This gradual shift of activity towards the left scalp region is consistent with the corresponding fNIRS results (Fig. 2A).

**Figure 3.**
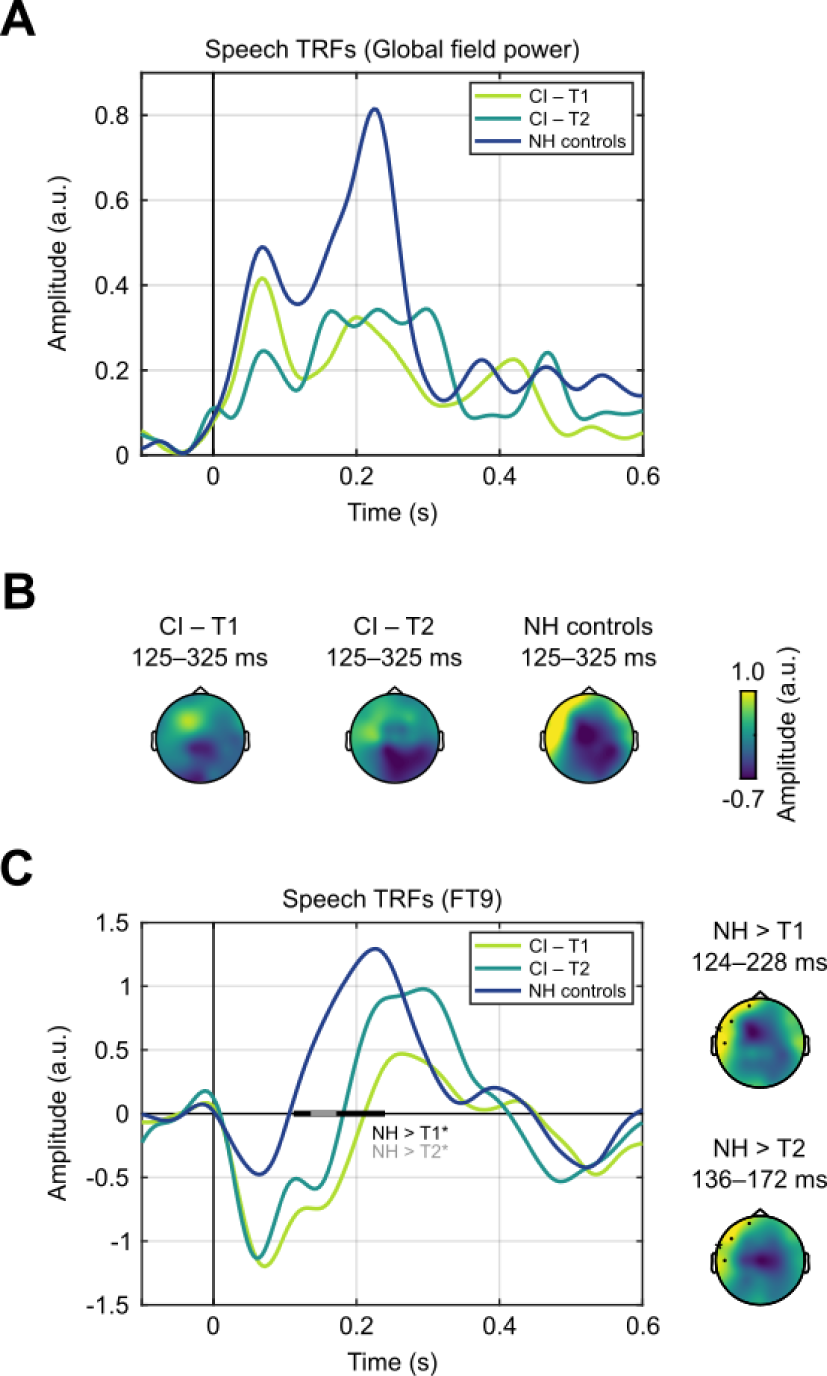
EEG results: speech experiment. A) Global field power of the speech envelope TRFs for each of the three experimental groups. B) Scalp maps showing the time-averaged TRF amplitudes from 125–325 ms for each group. C) TRFs for channel FT9, marked by a black asterisk in the scalp maps. The horizontal black and grey bars indicate time windows with significant amplitude differences between groups. In the corresponding scalp map on the rights, the time-averaged amplitude difference between groups is shown and significant channels are highlighted.

Differences of the speech envelope TRFs between groups were then statistically analysed via permutation-based (*n* = 10,000) one-sided independent-samples *t*-tests computed for each time point from 0–600 ms and each of the 29 electrodes. Significant inter-group differences were confined to the left fronto-temporal scalp region and the latency window of the second peak in the GFP waveforms. In this scalp region, this component had a positive polarity in all three groups, but amplitudes were larger and latencies shorter in the NH control group (Fig. 3C). Compared to the CI – T1 group, larger amplitudes for the NH controls were evident for 4 neighbouring electrodes from 124–228 ms (*p* < 0.05; averaged Cohen’s *d* = 1.03; upper scalp map in Fig. 3C). The same effect for the same cluster of electrodes was also observed for the comparison with the CI – T2 group, albeit with a shorter duration (*p* < 0.05, 136–172 ms, *d* = 0.63; lower scalp map in Fig. 3C). Despite a clear trend for larger responses in the CI – T2 group, no significant differences were found between the two CI groups with respect to this TRF component. These findings are also in line with the corresponding fNIRS results, where activity in left auditory areas was stronger in the NH control group compared to both CI groups, but no significant differences between the two CI groups were observed (Fig. 2C).

### 3.2. Vowel experiment

The fNIRS HbR topographies in response to the vowel sequences are shown in Fig. 4A, separately for each of the three groups. Same as in our previous study with adult CI users (Steinmetzger et al., 2022b), the auditory cortex activity elicited by these vowel stimuli was statistically evaluated by separately testing each channel included in two ROIs covering the STC in both hemispheres (left: ch# 5, 12, 14, & 16; right: ch# 27, 34, 36, & 38). The false discovery rate (FDR) across the ROI channels was controlled using the Benjamini-Hochberg procedure. For both the CI – T1 and CI – T2 groups, no significant activity was evident for any of the ROI channels. In contrast, significant bilateral activity in the posterior part of auditory cortex was observed in the NH control group (*p_(FDR)_* < 0.05, ch# 16 & 38, averaged Cohen’s *d* = 1.09).

**Figure 4.**
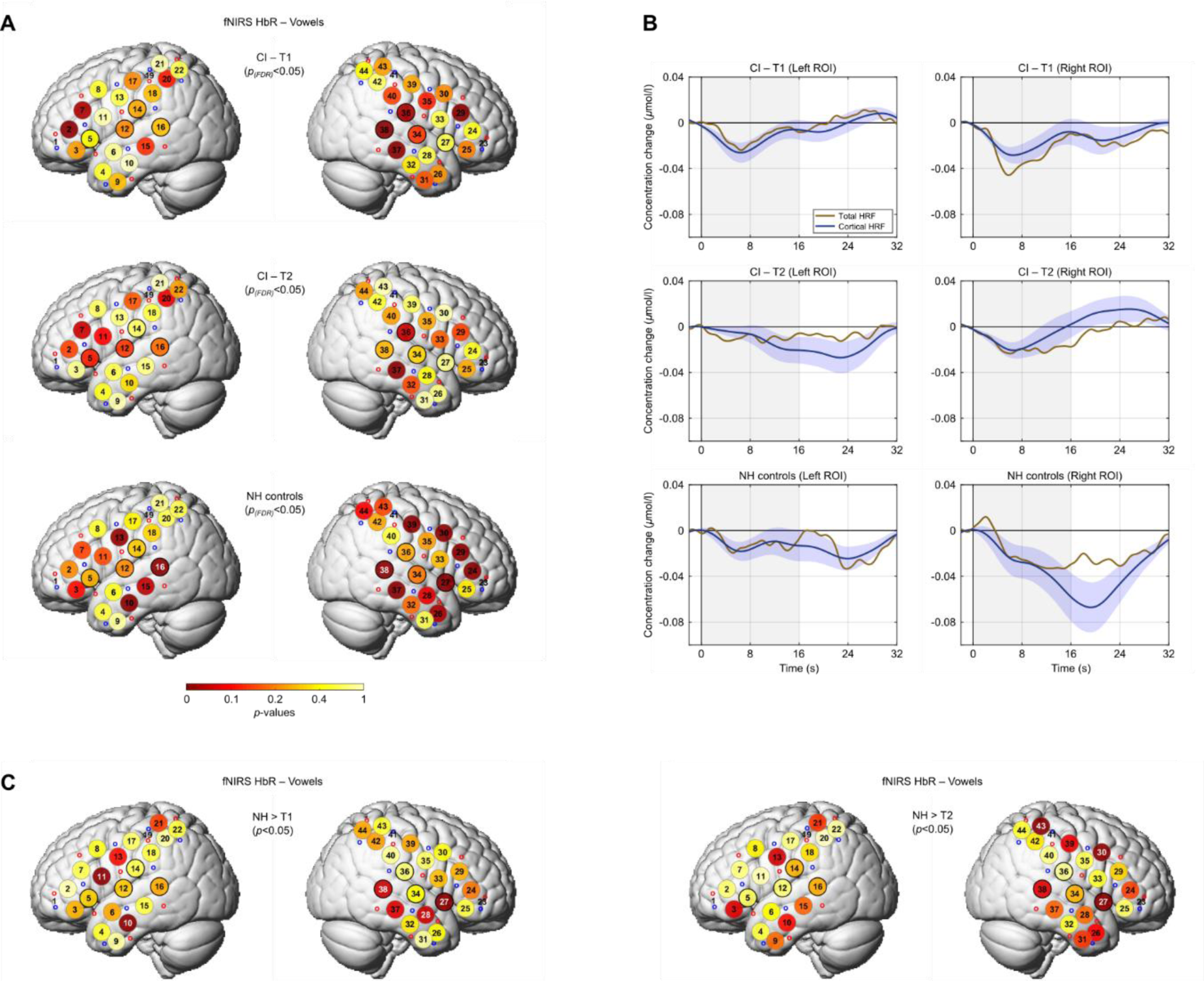
fNIRS results: vowel experiment. A) fNIRS HbR topographies of functional activity in response to vowel sequences for each of the three experimental groups. B) HRFs for each group and ROI. C) Comparison of the topographies of functional activity across groups. Apart from the specific ROI channels and the scaling of the HRFs, the details of the plot are the same as for Fig. 2.

The corresponding HRFs are shown in Fig. 4B, separately for each group and ROI. In agreement with the topographies, the amplitude changes evoked by the vowel sequences were very small in both CI groups, with peak amplitudes about three times smaller than in the preceding speech experiment. In the NH control group, however, the right ROI showed a pronounced response that was significantly larger compared to the left ROI (*t*_(11)_ = 2.57, *p* = 0.026, *d* = 0.74).

The functional activation patterns were then compared across groups (Fig. 4C), focussing on differences in the two auditory ROIs. For the NH group, activity for two channels in the right ROI was stronger than in the CI – T1 group (*p* < 0.05, ch# 27 & 38, *d* = 0.90). For the comparison with the CI – T2 group, significantly stronger activity was confined to the anterior portion of the right ROI (*p* < 0.05, ch# 27, *d* = 0.99). These results are thus consistent with our previous study (Steinmetzger et al., 2022b), where adult CI users with single-sided deafness also showed stronger activity along the right STC in response to these vowel sequences when listening via their NH ear compared to the CI ear. Other than in the previous paper, vowel sequences with VARIABLE compared to FIXED PROSODY did not evoke greater activity in any of the ROI channels for any of the groups (Suppl. Fig. 5), which may be due to the smaller sample sizes.

The concurrently recorded EEG data were analysed using typical ERP methodology, by averaging the responses to the individual vowels in the sequences. As can be seen in Fig. 5A, the vowel ERPs in the NH control group, but not the two CI groups, exhibited a pronounced P1 peak. Differences between the NH controls and the CI groups were confined to this time window. For the comparison with the CI – T1 group, a cluster-based permutation test (Maris & Oostenveld, 2007) returned a single highly significant cluster (∼56–200 ms, *t*_(cluster)_ = 1052.43, *p* < 0.001, *d* = 1.87). At its temporal midpoint of 128 ms after vowel onset, this cluster comprised 10 fronto-central channels (upper scalp map in Fig. 5A). The comparison with the CI – T2 group revealed a significant cluster with similar characteristics but a smaller temporal and spatial extent as well as a smaller effect size (∼68–196 ms, *t*_(cluster)_ = 456.82, *p* = 0.022, *d* = 1.33). Both tests were based on independent-samples *t*-tests for each time point from 0–800 ms with a cluster-forming threshold of *p* < 0.05 (one-sided), a minimum of 3 neighbouring electrodes per cluster, and 10,000 randomisations to determine the cluster *p*-values. Although the P1 effect relative to the NH controls was smaller for the CI – T2 group, there was again no significant difference between the two CI groups. Consistent with the fNIRS results, the ERP data hence showed that the vowel sequences elicited little auditory activity in the two CI groups, whereas large responses were observed in the NH controls.

**Figure 5.**
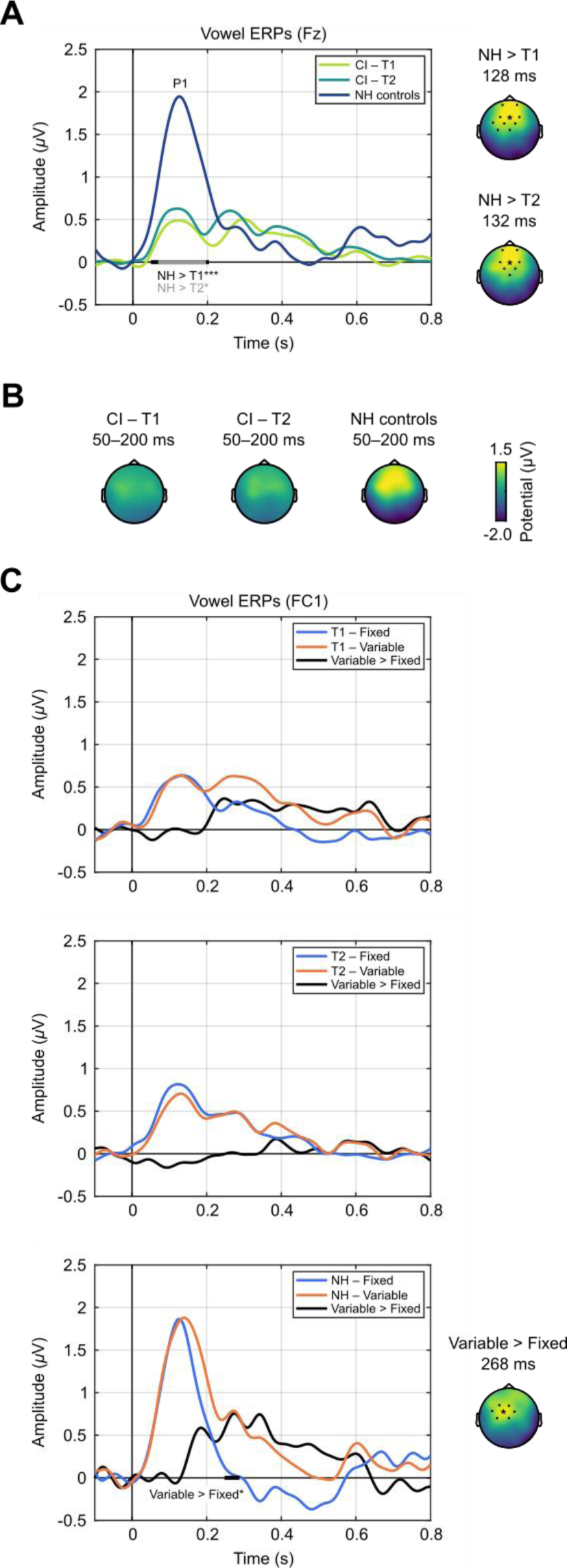
EEG results: vowel experiment. A) ERPs for channel Fz in response to vowel sequences for each of the three experimental groups. B) Scalp maps showing the time-averaged amplitudes from 50–200 ms. C) ERPs for the FIXED and VARIABLE PROSODY conditions, and the difference waveforms, for channel FC1. The details of the plot are the same as for Fig. 3.

To depict the distribution of the P1 component for the different groups, scalp maps showing the averaged amplitudes over a time window from 50–200 ms are displayed in Fig. 5B. Despite the substantially larger amplitudes in the NH control group, a similar fronto-central voltage distribution is evident in all three groups, implying that the underlying cortical sources were located in the auditory cortex in each case.

Finally, we examined whether the ERP amplitudes were larger in the VARIABLE PROSODY condition than in the FIXED PROSODY condition (Fig. 5C). Although there was a trend in this direction for the CI – T1 group, no such effect was evident for two CI groups when applying a cluster-based correction for multiple comparisons. For the NH control group, in contrast, a cluster consisting of 7 fronto-central electrodes reached significance (∼248–288 ms, *t*_(cluster)_ = 160.16, *p* = 0.048, *d* = 1.08), indicating that ERP amplitudes after the prominent P1 peak were enhanced in response to vowel sequences with VARIABLE PROSODY. Other than the fNIRS data, the current EEG data thus provided evidence that prosodic variations evoke greater auditory evoked responses, in line with previous data obtained from adult NH listeners (Steinmetzger et al., 2022a).

## 4. Discussion

### 4.1. Aberrant right-lateralised cortical activity in response to speech early after cochlear implantation

The fNIRS results of the speech experiment showed that cortical activity early after implantation (CI – T1 group) was strongest in right auditory areas, and that activity in this region was stronger compared to the more experienced CI users (CI – T2) and the NH controls. In contrast, speech-evoked auditory cortex activity in the latter two groups was more symmetrical across hemispheres, with a trend for a lateralisation to the left, and overall stronger responses for the NH controls. Hence, the current fNIRS results suggest a reduction of aberrant right-lateralised speech-evoked activity with more CI experience. The TRFs derived from the concurrently recorded EEG data revealed a similar pattern, as the scalp maps exhibited a topographical shift from the fronto-central scalp region in the CI – T1 group towards left fronto-temporal areas in the other two groups.

Previous ERP data obtained from pre- (Ni et al., 2021; Vavatzanidis et al., 2015; Vavatzanidis et al., 2018) and post-lingually deafened CI users (Sandmann et al., 2015) reported increased auditory evoked responses with more CI experience. However, these longitudinal studies used non-speech stimuli or isolated syllables rather than continuous speech, limiting comparability with the present findings. On the other hand, fNIRS studies that used running speech as stimulus yielded conflicting results and did not assess speech-evoked activity longitudinally. Wang et al. (2022) found no significant speech-evoked activity in either hemisphere for young pre-lingually deafened CI users directly after implantation and 1.5 months later, while Zhou et al. (2023) reported larger bilateral activity compared to a NH control group in experienced pre-lingually deafened children with CIs. Bilateral speech-evoked activity in auditory areas was also observed in experienced pre-lingually deafened CI users of the same age as in the current study (Mushtaq et al., 2020), consistent with the current CI – T2 results.

Furthermore, compared to NH controls, smaller bilateral responses for pre-lingually deafened CI users tested in adult age many years after implantation were observed (Levin et al., 2022), which is also in line with the present findings. The current results thus confirmed the absence of left-lateralised speech-evoked activity in more experienced pre-lingually deafened CI users. Yet, extending beyond previous studies, they also demonstrated that this bilateral activity pattern was preceded by abnormal right-lateralised activity in less experienced pre-lingually deafened CI users.

The lack of comparable longitudinal studies and the heterogeneity of the current sample of CI users make it particularly important to discuss possible confounding factors. Firstly, the deafness durations prior to implantation did not differ between the two CI groups (*p* = 0.732). Secondly, studies using non-speech stimuli such as pulse trains or tones have shown abnormally high contralateral cortical activity after prolonged unilateral deafness or late bilateral implantation in pre-lingually deafened paediatric CI users (Gordon et al., 2013; Polonenko et al., 2018) as well as animal models (Kral et al., 2013; Popescu & Polley, 2010; Tillein et al., 2016). However, only a small proportion of the datasets were obtained from subjects with unilateral left CIs in both groups (T1 = 6/17, T2 = 2/12). Therefore, it is unlikely that the right lateralisation of speech-evoked activity in the CI – T1 group is due to contralateral auditory input. Thirdly, although more datasets in the CI – T1 group were obtained from non-native German speakers (9/17 vs. 3/12), speech intelligibility scores were only slightly higher in the CI – T2 group (51.1 vs 61.7%). Thus, it also appears unlikely that the right lateralisation in the CI – T1 group was caused by a lower speech intelligibility.

Since the left lateralisation of speech processing is already apparent in NH new-borns and infants (Dehaene-Lambertz et al., 2002; Pena et al., 2003; H. Sato et al., 2012), long before the average age of the CI users tested here, the right lateralisation observed in the CI – T1 group seems to be a deafness-related effect. The results of the speech experiment hence imply that the language network in the left hemisphere needs time to develop following prolonged prelingual deafness. It is noteworthy in this context that only 4/13 and 9/13 children in the current sample received at least one CI within the proposed critical and sensitive time periods (∼3.5 & 7 years, respectively; Gilley et al., 2008; Sharma et al., 2002). Moreover, only 4/13 cases were bilaterally implanted.

Across both CI groups, the fNIRS topographies furthermore indicated less activity in the right hemisphere in case of higher speech intelligibility scores (Fig. 2E). This effect extended far beyond the auditory cortex and its spatial extent coincided with the right-hemispheric ventral attention network (VAN) comprising temporo-parietal junction, STG, as well as middle and inferior frontal gyri (Corbetta et al., 2008; Corbetta & Shulman, 2002). Deactivation of the VAN is associated with focussed attention, while transient VAN activity is thought to reflect a re-orientation to a salient auditory or visual stimulus (Kim, 2014; Larson & Lee, 2013). This finding might thus reflect a greater attentional focus on the speech stimuli in subjects with higher speech intelligibility scores.

### 4.2. EEG data reveal smaller and delayed speech-evoked activity in paediatric CI users

Previous EEG studies with CI users have consistently reported delayed ERP latencies and reduced amplitudes relative to NH control subjects, both in pre- (Gilley et al., 2008; A. Sharma et al., 2002) and post-lingually deafened participants (Kelly et al., 2005; Sandmann et al., 2015; Steinmetzger et al., 2022b; Viola et al., 2011). However, as discussed above, these findings were obtained via typical ERP methodology, in which short speech or non-speech stimuli were presented repeatedly to obtain averaged responses. Here, we showed that delayed and attenuated auditory evoked responses in CI users are also evident when using running, non-repetitive speech, the crucial stimulus for the evaluation of CI-based hearing.

In the NH control group, the morphology of TRFs resembled those in previous studies with NH adults, which also used a ridge regression approach to derive speech envelope-based TRFs from EEG data (Di Liberto et al., 2018; Fuglsang et al., 2017) or intracranial recordings (Golumbic et al., 2013). These studies found TRFs that exhibited a prominent positive peak with a latency of about 150–200 ms, framed by two somewhat smaller negative deflections. However, in contrast to those results, the scalp topography of the positive response component showed a pronounced left lateralisation for the NH controls, rather than a fronto-central distribution. This shift to the left temporal scalp region appears to be a child-specific effect, as recent data from 4-year-old NH children showed a similar distribution (Tan et al., 2022). Akin to the pattern observed in ERP studies with pre-lingually deafened CI users (Gilley et al., 2008; A. Sharma et al., 2002), the positive peak in the TRFs was delayed and had a smaller amplitude in the CI users, particularly those with less CI experience (Fig. 3A).

As in our previous study with adult CI users (Steinmetzger et al., 2022b), the current EEG data were only marginally affected by CI artefacts, which might be due to the use of continuous stimulation paradigms (Pantev et al., 2006). More generally, the fact that TRFs which revealed significant differences between CI users and NH controls could be derived from relatively short EEG recordings argues against claims that fNIRS is *per se* the most suitable technique to investigate cortical activity in CI users (Bortfeld, 2019; Pinti et al., 2018).

### 4.3. No evidence for right-lateralised processing of prosodic variations in paediatric CI users

For the NH controls, the fNIRS data showed right-lateralised responses to vowel sequences with prosodic variations, in agreement with our previous results obtained from NH adults (Steinmetzger et al., 2022a) and fNIRS data recorded from NH infants and children (Chen et al., 2022; Telkemeyer et al., 2011; Wartenburger et al., 2007) showing that prosodic variations elicit stronger activity in the right hemisphere. This lateralisation is thought to reflect that slow pitch changes, as found in speech and music, are preferentially processed by neural populations in the anterior part of the right superior temporal cortex (Johnsrude et al., 2000; Patterson et al., 2002; Zatorre & Belin, 2001). Although there was a small trend in this direction, no such effect was observed in both CI groups and no differences between the FIXED and VARIABLE PROSODY conditions were evident. This finding is in line with two recent cross-sectional fNIRS studies (Chen et al., 2022; Wang et al., 2021) that also did not report right-lateralised processing of prosodic features in paediatric CI users. In addition to those two studies, the current results showed that the processing of prosodic information did not change with more CI experience. The corresponding ERP data revealed a similar picture, with much larger P1 amplitudes in the NH control group and no differences between the two CI groups. Moreover, larger responses in the VARIABLE PROSODY condition were only evident for the NH controls.

Prosodic variations are difficult to perceive with current CI systems, as pitch information is primarily transmitted via weak temporal envelope cues (Green et al., 2005; Nakata et al., 2012; Steinmetzger & Rosen, 2018). Yet, our previous fNIRS and EEG data provided clear evidence for right-lateralised processing of prosodic variations in post-lingually deafened adult CI users (Steinmetzger et al., 2022b). This suggests that prior experience with normal acoustic hearing might be required to perceive these stimulus features. However, it should also be noted that both in the present experiment and the two other fNIRS studies that investigated the cortical processing of prosody in paediatric CI users (Chen et al., 2022; Wang et al., 2021), response amplitudes were very small in comparison to the respective NH control groups. The low signal-to-noise ratio of the data therefore made it difficult to reveal differences between experimental conditions as well as the two CI groups. For example, the comparison of the ERPs in the FIXED and VARIABLE PROSODY conditions in the CI – T1 group revealed a clear trend for more activity in the latter condition (Fig. 5C), but due to the small overall ERP amplitudes this did not result in a significant difference.

Related to this, the present fNIRS data showed that auditory cortex activity in response to running speech was about three times larger compared to the proceeding vowel experiment, despite a much lower number of trials. At the same time, the fNIRS amplitudes in the vowel experiment were largely similar in size to those found in our previous studies with adult NH listeners (Steinmetzger et al., 2022a) and CI users (Steinmetzger et al., 2022b) in which the same stimuli were used. A similar comparison across studies is not possible based on EEG data as the morphologies of auditory ERPs recorded from children and adults differ markedly (Ponton et al., 1996; Ponton et al., 2002), which constitutes a major advantage in favour of BOLD-based measures such as fNIRS.

Moreover, since MRI analyses indicated only minor structural differences between the brains of adults and children of the age tested here (Fonov et al., 2011), the same fNIRS probe layout and ROIs as in the previous studies were used. Although the scalp-brain distance is somewhat smaller in children and the infrared light hence reaches a little deeper into the cortex (Beauchamp et al., 2011), the vowel-evoked activity should thus be readily comparable across these studies. The present fNIRS data thus clearly demonstrate that continuous speech is better suited to evoke large auditory cortex responses in paediatric subjects than artificially created stimuli such as vowel sequences.

## Data and code availability

Stimuli, fNIRS and EEG data, as well as the code used to process the data are all available at: https://doi.org/10.17605/osf.io/2sx69.

## Author contributions

B.K.: Formal analysis, Investigation, Data Curation, Writing – Original Draft. M.S.R.: Investigation, Resources. A.G.: Methodology, Software, Writing – Review & Editing. M.A.: Conceptualization, Writing – Review & Editing, Supervision, Funding acquisition. M.P.: Conceptualization, Resources, Supervision, Project administration, Funding acquisition. A.R.: Conceptualization, Resources, Writing – Review & Editing, Supervision, Project administration, Funding acquisition. K.S.: Conceptualization, Methodology, Software, Formal analysis, Investigation, Data Curation, Writing – Original Draft, Visualization, Supervision, Funding acquisition

## Acknowledgements

We would like to thank Elisabeth Munk for helping with the recruitment of the participants and Anne van der Kant for suggesting the use of a car seat for testing the children.

## Supplementary data

Supplementary material for this article is available with the online version here: *<u>paste link here</u>*.

**Supplementary Figure 1.**
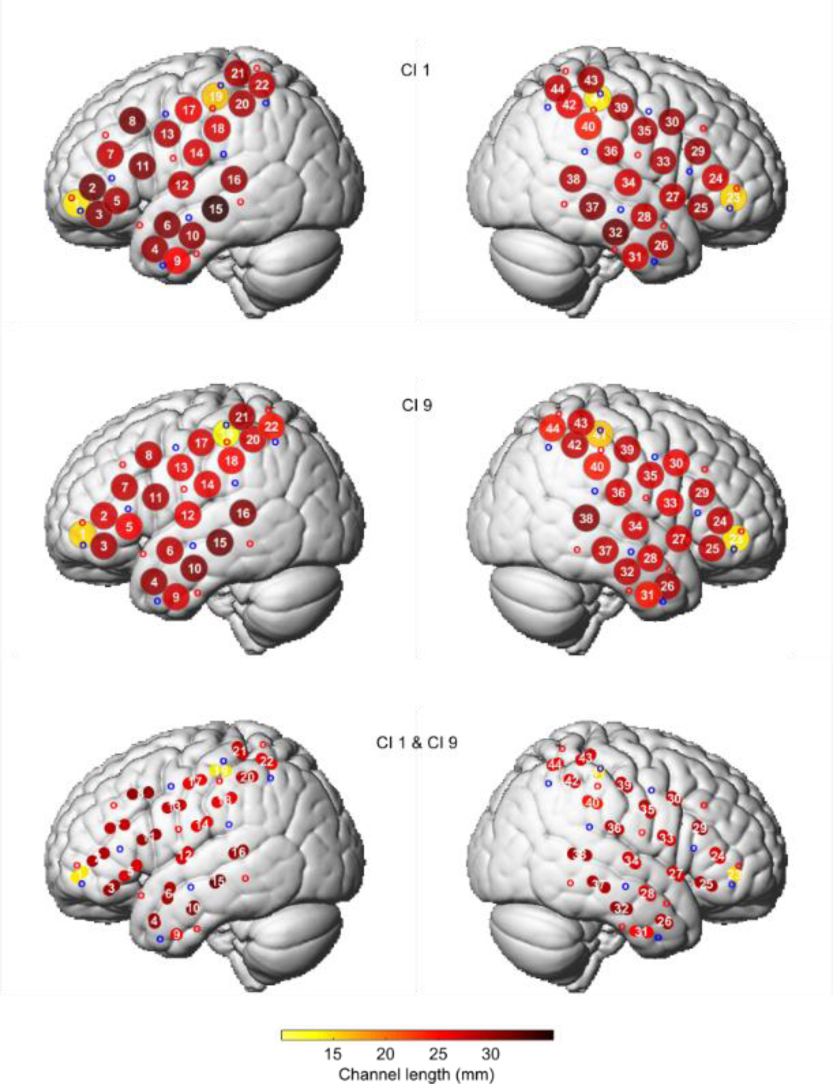
fNIRS measurement layout. The positions of the source and detector optodes (small red and blue circles, respectively) as well as the resulting measurement channels (white numbers) for subjects CI 1 and CI 9, rendered onto the ICBM-152 cortical template surface. The colour of the disc at the respective channel location indicates the channel length. In the bottom row, the digitisations of both subjects were averaged. Here, the points surrounding the channel numbers indicate the positions of the individual subjects.

**Supplementary Figure 2.**
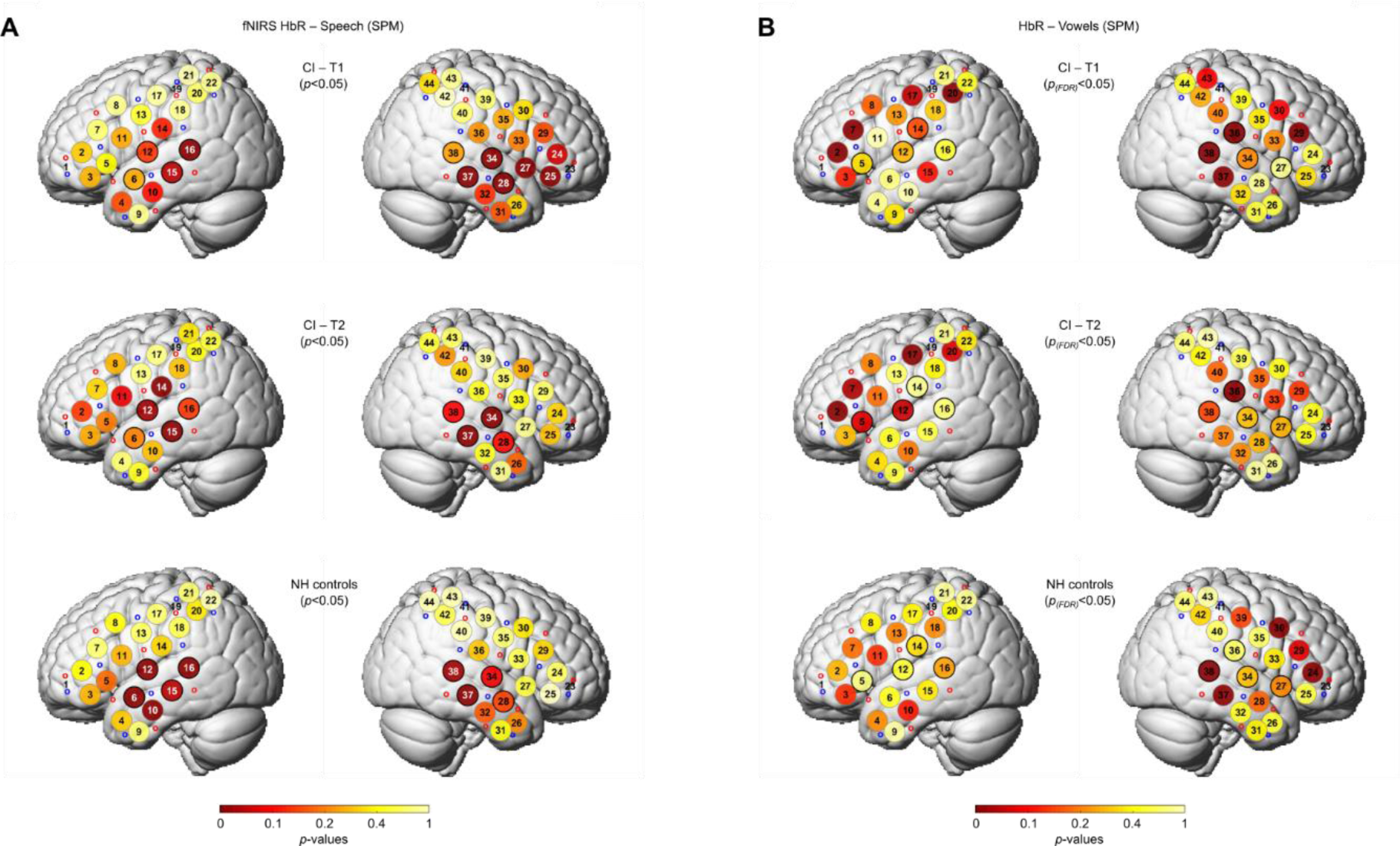
fNIRS SPM results. fNIRS HbR topographies of functional activity in response to speech (A) and vowel sequences (B) for each of the three experimental groups. The details of the plot are the same as for *Figs. 2A and 4A*, except that the results are based on SPM-based modelling of the responses rather than mean amplitude measures.

**Supplementary Figure 3.**
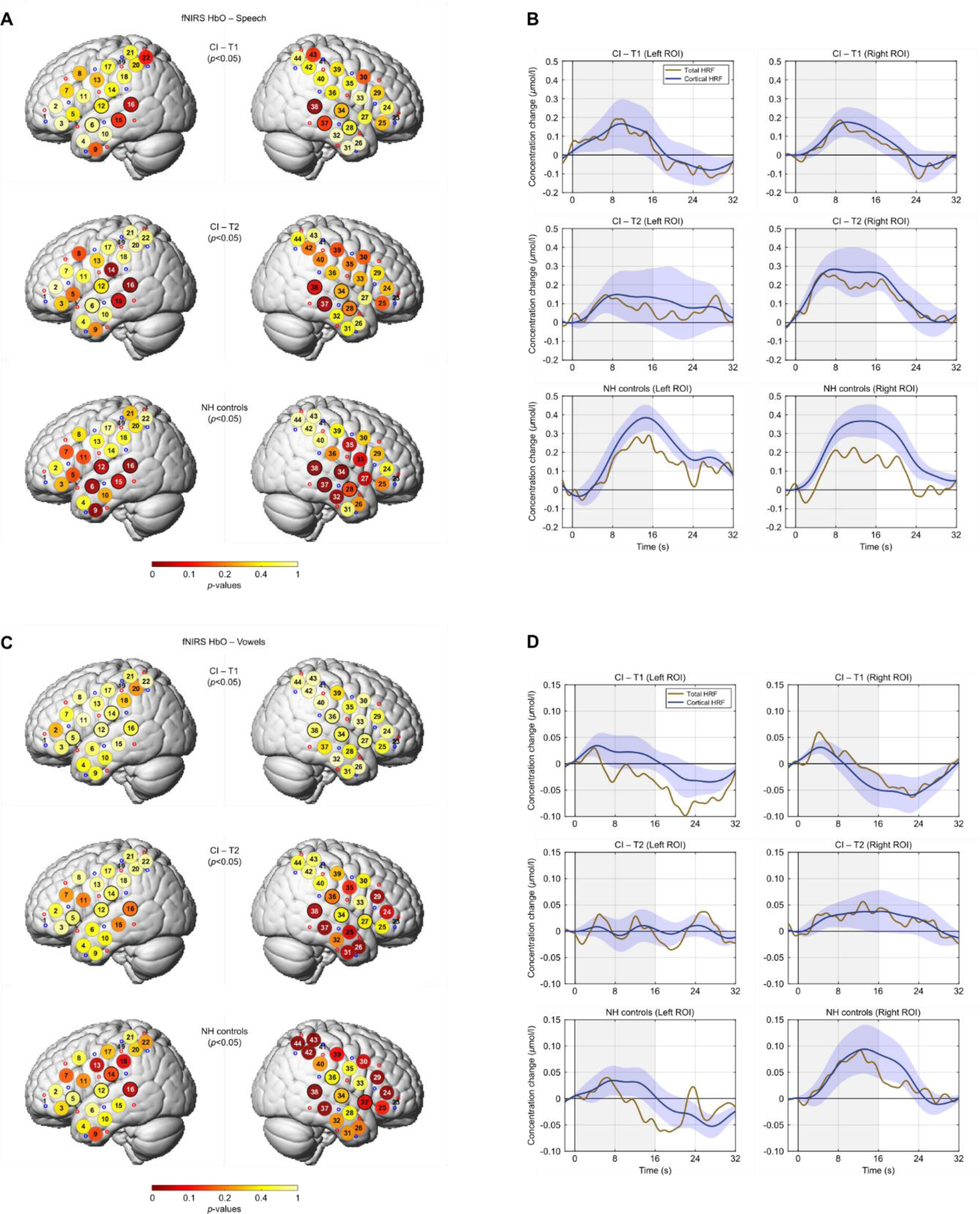
fNIRS HbO results. fNIRS HbO topographies of functional activity in response to speech (A) and vowel sequences (B) for each of the three experimental groups. The details of the plot are the same as for *Figs. 2A* & B, except that the results are based on HbO instead of HbR data.

**Supplementary Figure 4.**
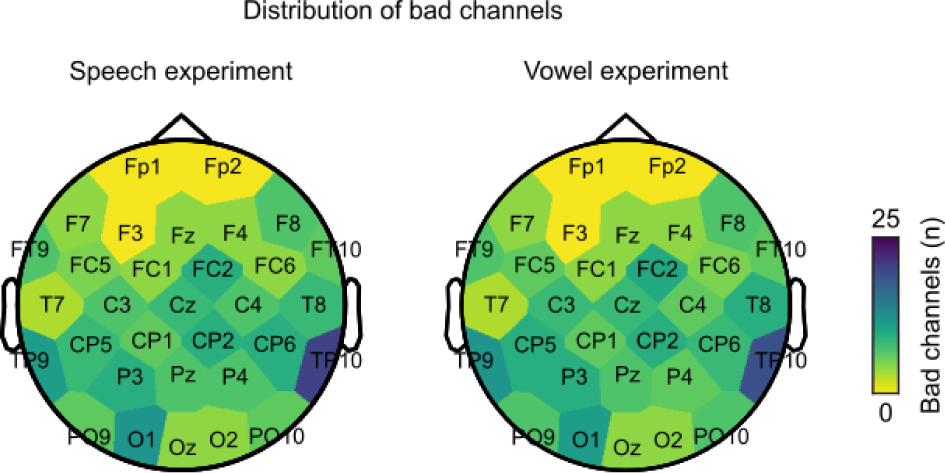
Scalp maps of bad EEG channel counts in the speech and vowel experiments.

**Supplementary Figure 5.**
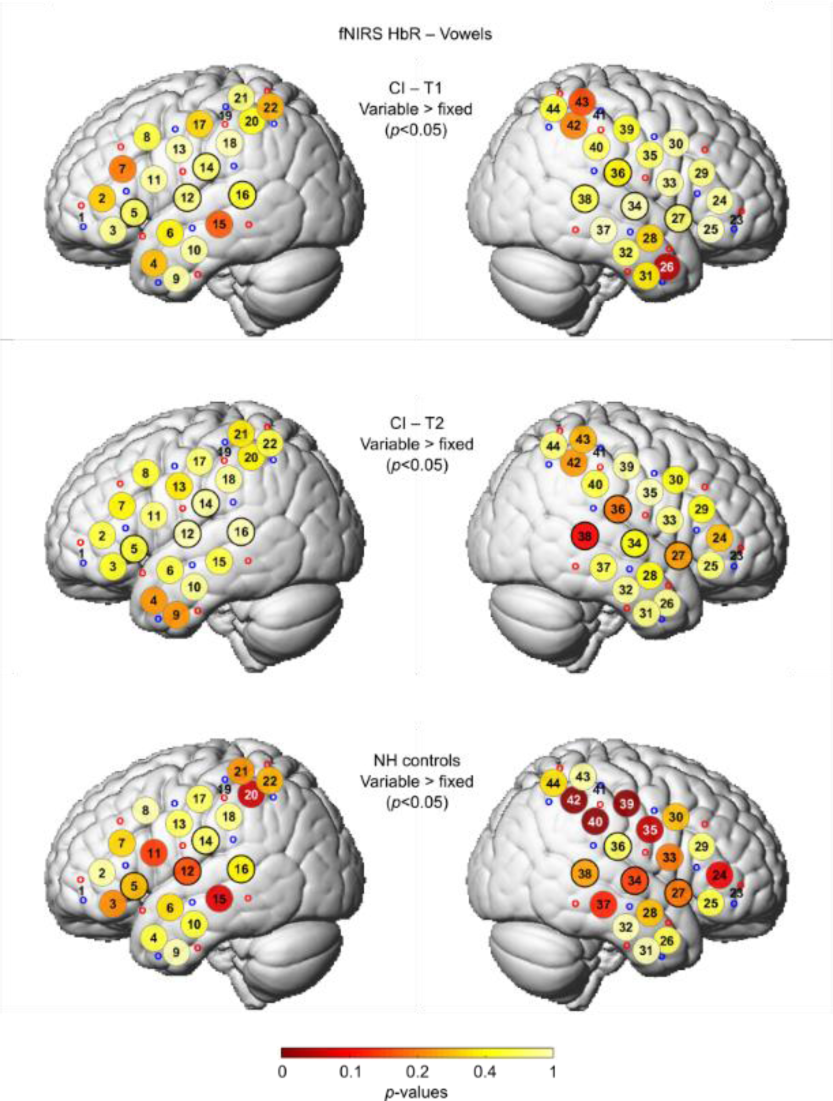
fNIRS results: vowel experiment. fNIRS HbR topographies of functional activity showing where the cortical activity in the VARIABLE PROSODY condition exceeded that in the FIXED PROSODY condition, separately for each of the three experimental groups.

## Abbreviations

CI: Cochlear implant
EEG: Electroencephalography
ERPs: Event-related potentials
FDR: False discovery rate
fNIRS: Functional near-infrared spectroscopy
GLM: General linear model
HbO: Oxygenated haemoglobin
HbR: De-oxygenated haemoglobin
HRF: Haemodynamic response function
NH: Normal-hearing
PET: Positron emission tomography
ROI: Region of interest
SACF: Summary autocorrelation function
SCI: Scalp coupling index
SP: Sustained potential
SPM: Statistical parametric mapping
STC: Superior temporal cortex
STG: Superior temporal gyrus
STS: Superior temporal sulcus
TRF: Temporal response function
VAN: Ventral attention network

